# Dynamic functional connectivity markers of objective trait mindfulness

**DOI:** 10.1101/287540

**Authors:** Julian Lim, James Teng, Amiya Patanaik, Jesisca Tandi, Stijn A. A. Massar

**Affiliations:** Center for Cognitive Neuroscience, Neurosciences and Behavioral Disorders Department, Duke-NUS Medical School. Singapore 169857

**Keywords:** mindfulness, resting-state fMRI, dynamic functional connectivity, state transitions, breath-counting

## Abstract

While mindfulness is commonly viewed as a skill to be cultivated through practice, untrained individuals can also vary widely in dispositional mindfulness. Prior research has identified static neural connectivity correlates of this trait. Here, we use dynamic functional connectivity (DFC) analysis of resting-state fMRI to study time-varying connectivity patterns associated with naturally varying and objectively measured trait mindfulness. Participants were selected from the top and bottom tertiles of performers on a breath-counting task to form high trait mindfulness (HTM; N = 21) and low trait mindfulness (LTM; N = 18) groups. DFC analysis of resting state fMRI data revealed that the HTM group spent significantly more time in a brain state associated with task-readiness – a state characterized by high within-network connectivity and greater anti-correlations between task-positive networks and the default-mode network (DMN). The HTM group transitioned between brain states more frequently, but the dwell time in each episode of the task-ready state was equivalent between groups. These results persisted even after controlling for vigilance. Across individuals, certain connectivity metrics were weakly correlated with self-reported mindfulness as measured by the Five Facet Mindfulness Questionnaire, though these did not survive multiple comparisons correction. In the static connectivity maps, HTM individuals had greater within-network connectivity in the DMN and the salience network, and greater anti-correlations between the DMN and task-positive networks. In sum, DFC features robustly distinguish HTM and LTM individuals, and may be useful biological markers for the measurement of dispositional mindfulness.

## 1. Introduction

The last decade has seen a surge of interest in the cognitive neuroscience of mindfulness (Tang, et al., 2015), driven largely by growing evidence of its benefits for cognition, health, and well-being (Chiesa and Serretti, 2009; Chisea, et al., 2011; Kabat-Zinn, 1991; Sedlmeier, et al., 2012). Mindfulness is a state characterized by awareness and attention to present-moment thoughts and sensations while adopting an accepting, non-judgmental stance towards those experiences. While the preponderance of research has focused on cultivating mindfulness through formal programs or informal exercises, a relatively understudied topic is the biological and psychological underpinnings of trait mindfulness in those not exposed to mindfulness training. While intervention studies can provide insights into these trait differences, they are also susceptible to many cognitive, social, and cultural confounds, which might make interpretation difficult. As an alternative, understanding the natural variation of trait mindfulness may provide a cleaner picture of how it is instantiated in the human brain.

To date, neuroimaging studies have implicated regions across multiple brain networks associated with the practice of mindfulness. Within the salience network, the insula and the anterior cingulate are key nodes that consistently increase in volume, activation, and functional connectivity with meditation practice (Farb, et al., 2013; Holzel, et al., 2008; Holzel, et al., 2007; Luders, et al., 2012; Lutz, et al., 2013; Tang, et al., 2010). Numerous studies have also shown reduced default mode network (DMN) activation and stronger coupling after mindfulness training (Berkovich-Ohana, et al., 2016; Brewer, et al., 2011; Creswell, et al., 2016), particularly in posterior cingulate cortex and the precuneus. Finally, connectivity between the DMN and salience (or ventral attention) network in particular may play an important role in supporting mindfulness (Doll, et al., 2015).

Relatively few studies have investigated whole-brain functional connectivity in the context of mindfulness training. A study by Kilpatrick and colleague using independent components analysis showed altered connectivity following a mindfulness-based stress reduction course that was mostly localized to sensory cortex and the salience network (Kilpatrick, et al., 2011). Tang et al. (2017) used multi-voxel pattern analysis to classify resting state fMRI (rs-fMRI) scans collected before and after a two-week body-mind training course, and found increased functional connectivity distributed widely across the cortex, particularly in frontal, temporal, and occipital nodes.

Recently, dynamic functional connectivity (DFC) has emerged as a promising tool for the analysis of rs-fMRI data (Handwerker, et al., 2012; Hutchison, et al., 2013). This method commonly measures the covariance of BOLD signal across regions of interest (ROIs) in sliding time windows of a fixed interval (Allen, et al., 2014). These patterns can then be clustered into a pre-specified number of dynamic connectivity states (DCS) that can be robustly recovered across datasets and analysis methods (Abrol et al., 2017). Given that the brain is by nature a dynamic system (Tognoli and Kelso, 2014), DFC may capture functionally important facets of time-varying connectivity that might be missed by aggregate or static measures.

Recently, Mooneyham et al. (2017) used DFC analysis and identified a brain state correlated with dispositional mindfulness: this “focused attention” state featured strong within-network connectivity in the salience and executive control network, and reduced connectivity between these task-positive networks and the DMN. This segregated brain configuration in the resting state can be thought of as “task-readiness”, as smaller updates are necessary when task demands are imposed (Schultz and Cole, 2016).

In the present study, we selected good and poor performers from a large sample (N = 125) of participants who had previously performed a breath counting task (BCT) (Wong, et al., in press), a reliable, valid, and objective measure of trait mindfulness (Levinson, et al., 2014). We labeled these groups high and low trait mindfulness respectively (HTM/LTM). These participants underwent rs-fMRI scanning, and our specific aim was to identify differences in static and dynamic functional connectivity patterns in these data. As predicted, individuals high on trait mindfulness switched more often between DCS, and spent more time overall in a state associated with task-readiness, and static connectivity differed between the groups in several of the *a priori* networks identified. These findings shed some light on the connectivity metrics associated with the mindful brain.

## 2. Materials and Methods

### 2.1 Study procedure

Participants in this experiment were selected from a larger group of subjects (N = 125) who previously completed the Breath Counting Task (BCT) (see Section 2.2 and (Wong, et al., in press)). Participants were classified as having high trait mindfulness (HTM) if they were in the top tertile of BCT performers, and as having low trait mindfulness (LTM) if they were in the bottom tertile, based on overall task accuracy. In this first session, participants also underwent a 20-minute version of the Psychomotor Vigilance Test (PVT) (Dinges, 1995).

HTM and LTM participants were invited to return to the lab for fMRI scanning. To control for time-of-day effects, all scans took place between 1400 h and 1700 h. We were able to obtain data from 21 HTM (mean (SD) age = 23.7 (3.4); 8 male) and 18 LTM (mean (SD) age = 21.9 (2.3); 5 male) individuals. During this second testing session, participants completed a second trial of the BCT before undergoing the fMRI procedure described below. Following the fMRI scan, they completed the Five-Facet Mindfulness Questionnaire (FFMQ) (Baer, et al., 2008), a self-report measurement of multiple facets of trait mindfulness. Participants in the second session were compensated S$25.

This study was approved by the National University of Singapore Institutional Review Board and all participants provided written informed consent.

### 2.2 Breath-counting task

Trait mindfulness was assessed objectively via a 20-minute Breath Counting Task (BCT; (Levinson, et al., 2014)). In this task, participants were instructed to silently count their breaths from 1 to 9 repeatedly, to indicate breaths 1-8 with by pressing the left arrow key, and breath 9 with the right arrow key on a standard QWERTY keyboard. They were instructed to press the spacebar if they noticed that had lost count. We considered a right-arrow-key press or spacebar press to indicate the termination of a cycle. Correctly completed cycles were those composed of 8 left-arrow-key presses followed by a right-arrow-key press. BCT accuracy was calculated by dividing the number of correctly completed cycles by the total number of cycles.

### 2.3 Psychomotor Vigilance Test

The PVT is a reaction time (RT) test that is a sensitive assay of sustained attention (Dinges, 1995). In the task, participants monitor a rectangular box in the center of a screen and respond as quickly as possible to the appearance of a millisecond counter. The number of lapses (RT > 500 ms) on this test is a robust marker of vigilance (Basner and Dinges, 2011). PVT stimuli were presented using Psychtoolbox (Brainard, 1997) in MATLAB 2012a.

### 2.4 fMRI scans

Functional MRI scans were collected on a 3T Siemens Prisma system (Siemens, Erlangen, Germany). Two functional runs were collected: one 8-minute 20-second (250 TRs) eyes-open resting state (RS) scan, and one run of a finger-tapping task of equivalent length. Data from the task-based scan are not reported in this manuscript. We used a gradient echo-planar imaging sequence (TR = 2000 ms, TE = 30 ms, FA = 90°, FoV = 192 × 192 mm, voxel size = 3 × 3 × 3 mm) to acquire functional data. Following the functional scans, high-resolution structural images were collected with an MPRAGE sequence (TR = 2300 ms, TI = 900 ms, FA = 8°, voxel dimension = 1×1×1 mm, FOV = 256 × 240 mm). Images were preprocessed following the procedure detailed in Yeo et al., (2015) which included discarding the first 2 volumes, slice-time correction, head-motion correction, structure-function data alignment, linear trend removal, and low-pass temporal filtering. Linear regression was performed to remove effects associated with head motion, white matter, ventricular, and global signal. As we intended to perform dynamic functional connectivity analysis on the resultant time courses, motion scrubbing was not conducted. Cortical ROIs were delineated using the 17-network (114 ROI) cortical parcellation scheme of Yeo et al. (2011) (see Supplementary Table 1 for full list of ROIs), and time courses were averaged across each of the ROIs before being entered into the connectivity analysis.

### 2.5 rs-fMRI analysis

DFC analysis is a technique that is used to evaluate time-varying fluctuations of connectivity patterns in the brain. The typical output of this analysis is a series of connectivity matrices that can then be assigned (either discretely or probabilistically) to dynamic connectivity states (DCS). These states are thought to represent metastable brain configurations that may support different modes of information transfer. While a diversity of methods exists for calculating DFC, all of these require data to be transformed prior to estimating connectivity, and for a relational function between ROI/node pairs to be specified (Thompson and Fransson, 2017)

We carried out DFC analysis on our rs-fMRI data using the multiplication of temporal derivatives (MTD) procedure described by Shine et al. (2015). This method entails estimating the coupling between each pairwise set of ROIs by obtaining the normalized product of their first-derivative time courses. As recommended, we estimated connectivity at each time point by computing a simple moving average of the MTD time course using a window size of 7 TRs, yielding a total of 240 coupling matrices per subject. Each of these windows contained 6441 ((114 × 113)/2) unique coupling values. We selected this method as we were particularly interested in detecting state transitions, and the MTD procedure is more sensitive than traditional sliding window correlation methods in detecting these (Shine, et al., 2015).

Following this procedure, averaged MTD values were then concatenated across all participants, and k-means clustering was performed to classify each matrix using squared Euclidean distance as the cost function. For each subject, the proportion of windows classified into each state was then computed. We also extracted the number of state transitions observed during the run for each participant, and the average dwell time in each episode of a consistently obtained “task-ready” brain state (see Section 3.2).

### 2.6 Experimental design and statistical analysis

The threshold of statistical significance for all analyses was set at α = 0.05. Dependent variables of interest from the DFC analysis were compared between the HTM and LTM groups using independent-samples t-tests. These included: proportion of time spent in dynamic connectivity states, number of state transitions, and average dwell time. This process was repeated for a range of clusters from k = 3 to k = 7. To control for arousal, we conducted a secondary analysis where the number of PVT lapses was entered into the model as a between-subjects covariate. Pearson’s correlation was used to assess the linear relationship between time spent in the task-ready brain state and total score on the FFMQ.

Additionally, we obtained an estimate of static functional connectivity (SFC) by averaging the windowed MTD data for each subject. Mean connectivity within and between several networks of interest (all ROIs within the default mode, dorsal attention (DAN), ventral attention (VAN) and executive control (ECN) networks) was computed. From here, we refer to the DAN, VAN and ECN collectively as “task-positive” networks. The parcellation scheme and list of ROIs within these networks is available at https://surfer.nmr.mgh.harvard.edu/fswiki/CorticalParcellation_Yeo2011. We then conducted independent samples t-tests to assess if these SFC variables differed between groups. False discovery rate (FDR) correction using the method of Benjamini and Hochberg (1995) was applied across these tests to adjust for multiple comparisons.

Analysis of resting-state fMRI was performed using publicly available and in-house scripts in MATLAB 2012a (Mathworks Inc., Natick, MA, USA), and statistical analysis was conducted on extracted variables using SPSS version 24 (Armonk, NY: IBM Corp).

## 3. Results

### 3.1 Breath counting performance is reliable over time

Participants performed the BCT on two separate occasions, and were selected for fMRI scanning based on their session 1 performance. In session 1, the HTM group had a mean (SD) accuracy of 90.2 (5.5)%, and the LTM group had a mean (SD) accuracy of 56.1 (9.2)%. Performance differed significantly between the two groups (t_37_ = 12.11, p < 10^−13^). In session 2, we observed some regression to the mean, with 88.2 (11.9)% accuracy in HTM, and 73.5 (16.6)% accuracy in LTM. However, session 2 accuracy was still significantly different between the two groups (t_37_ = 3.20, p = .003). The two-way mixed intra-class correlation of BCT accuracy between sessions 1 and 2 was 0.48 (p = .001)

### 3.2 Dynamic functional connectivity analysis reveals two reproducible states

We performed DFC analysis on the rs-fMRI data using the MTD approach (Shine, et al., 2015) followed by k-means clustering on the resultant MTD windows pooled across all participants. We performed this clustering using values of k ranging from 3 to 7, and across all of these solutions found two consistently reproducible brain states. These polar states resembled those previously reported by Wang et al. (2016) despite differences in the method used in this study to analyze DFC. The first, and less common state (State 1: 25.1% of windows in the k = 3 solution) features strong within-network correlations in the DMN and the ventral attention/salience network (VAN), and larger anti-correlations between task-positive networks and the DMN. The second, more common state (State 2; occurring in 74.8% of the windows in the k = 3 solution) is characterized by lower within-network correlations and relatively small anti-correlations between task-positive networks and the default mode network (DMN). We call these the “task-ready” and “idling” states respectively. The centroids of the two states and a contrast map between them are shown in Figure 1.

**Figure 1:**
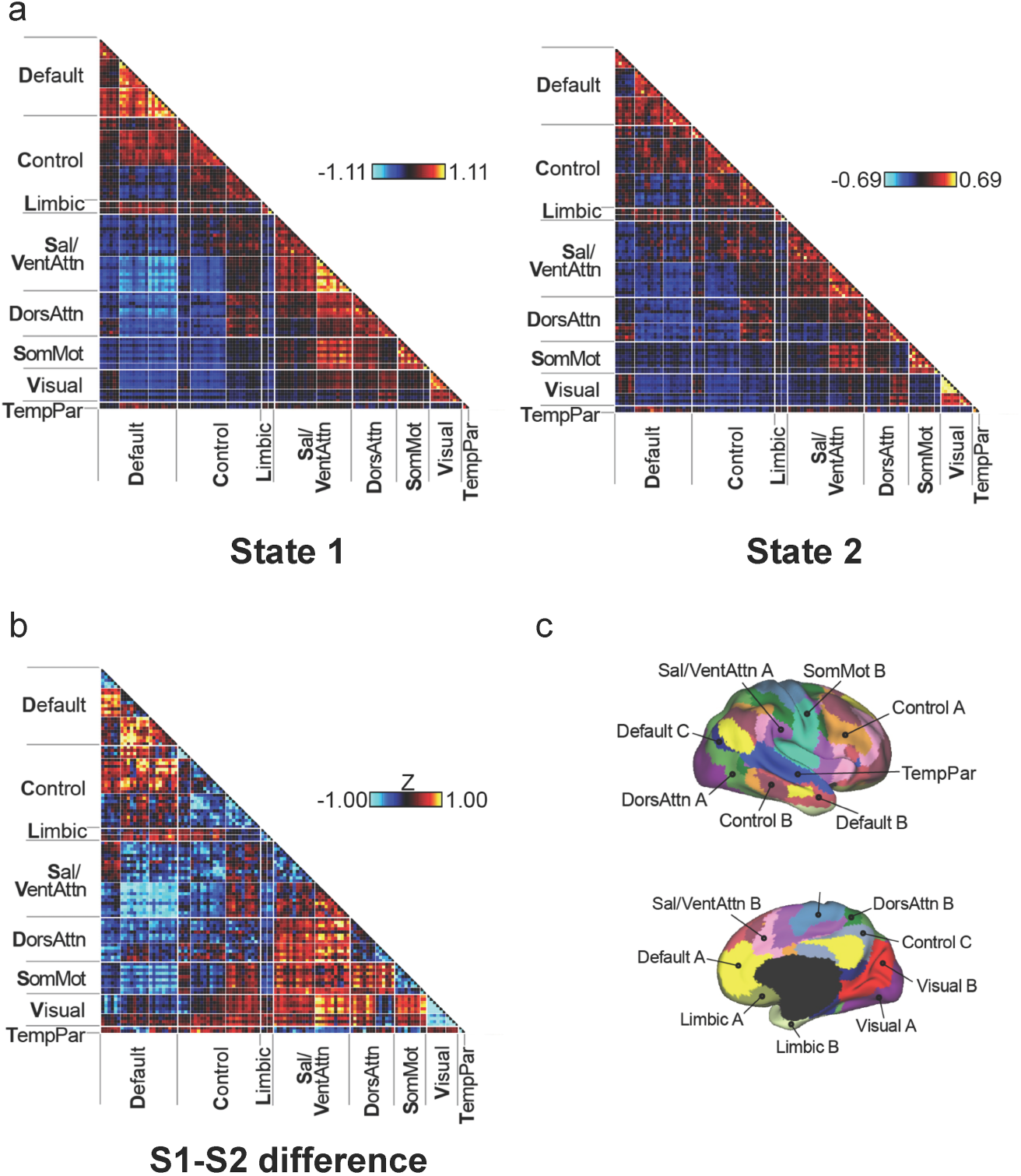
Dynamic connectivity states in resting-state fMRI. a) Representative (centroid) connectivity matrices of State 1 and State 2. Each row and column in the matrix represents one node (ROI), with individual cells representing connectivity between pairs of nodes. Within network comparisons are along the diagonal of the matrix (triangles), and between-network comparisons are the quadrilaterals bounded by the thicker white lines. State 1 is a “task-ready” state with high within-network connectivity and greater anti-correlations between the default-mode network and task-positive networks. State 2 is an “idling” state with relatively higher between-network connectivity and lower within-network connectivity. Units of connectivity (shown in the colour bars) are average coupling scores derived from the multiplication of temporal derivatives method. b) Z-scored difference between the State 1 and State 2 centroids. Connectivity differences between these centroids are greatest in nodes within the default mode network and the salience/ventral attention network. c) The parcellation scheme of Yeo et al. (2011) used to derive the 114 cortical ROIs. DorsAttn = dorsal attention network; Sal/VentAttn = salience/ventral attention network

We note that the third brain “state” in the k = 3 solution captured signal dropout over a short 8-TR period attributable to head motion in a single subject. Censoring these windows prior to k-means clustering removes this cluster while essentially preserving the results presented here. From here on, we focus on the results from the k = 3 solution for our primary analysis, since this is the lowest-dimensional solution at which our states of interest appear. Results from the higher-dimensional solutions and weighted averages are also included for completeness.

### 3.3 High trait individuals spend more time in the task-ready state

We next tested whether HTM and LTM participants spent different amounts of time in the two reproducible brain states. In the k = 3 solution, HTM participants spent significantly more time in State 1 (t_37_ = 2.29, p = .03) and less time in State 2 (t_37_ = 2.26, p = .03) (Figure 2). These differences were largely preserved as we increased the number of clusters allowed in the k-means analysis (Table 1). As the percentage of time spent in State 2 did not differ significantly across all clusters, we calculated a weighted effect size of these differences; this composite measure was not statistically significant (Cohen’s d = 0.63, p = .07).

**Table 1.**
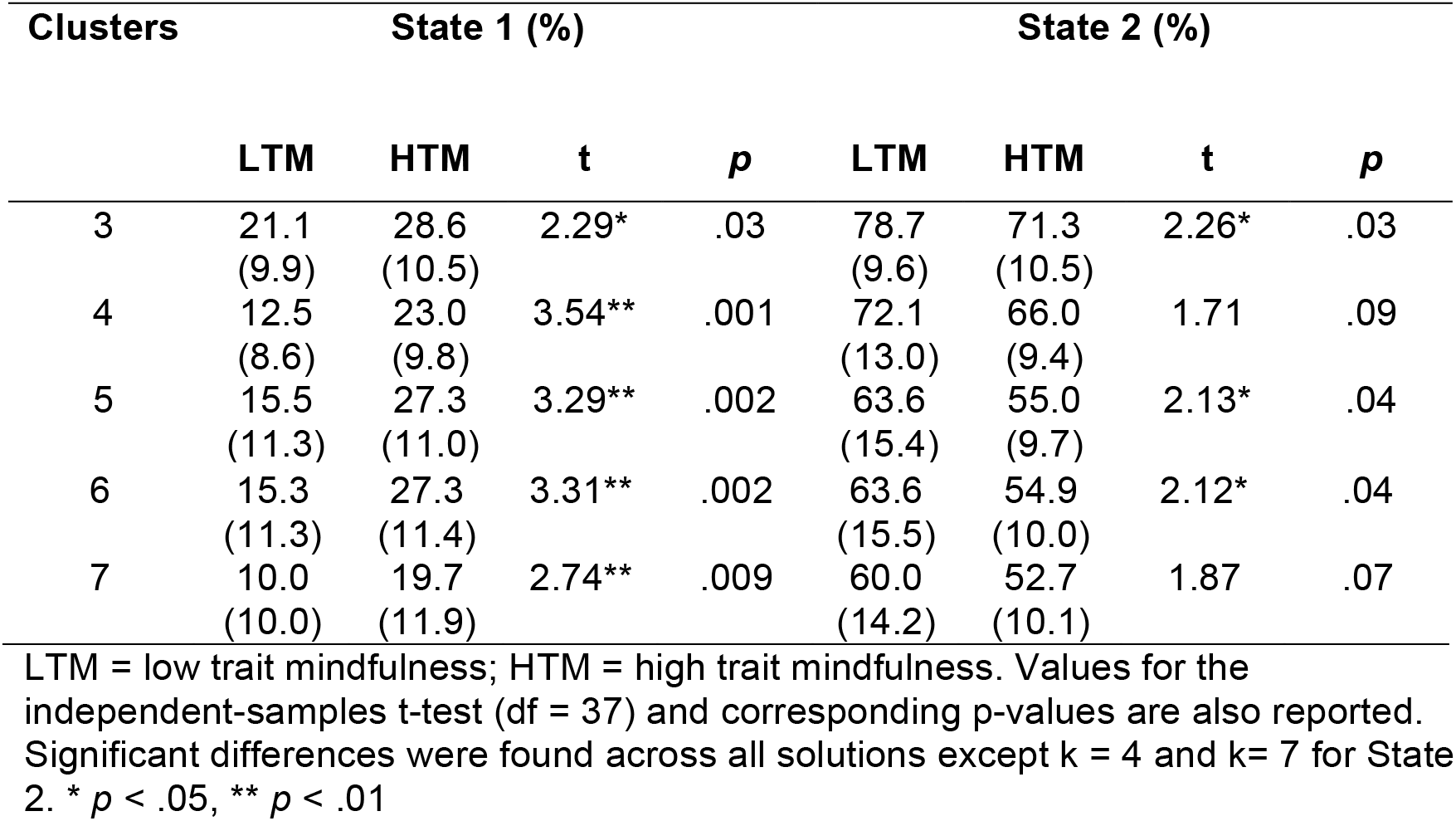
Mean and standard deviation of the percentage of time spent in States 1 and 2 for increasing values of k.

| Clusters | State 1 (%) |  |  |  | State 2 (%) |  |  |  |
| --- | --- | --- | --- | --- | --- | --- | --- | --- |
|  | LTM | HTM | t | p | LTM | HTM | t | p |
| 3 | 21.1<br>(9.9) | 28.6<br>(10.5) | 2.29* | .03 | 78.7<br>(9.6) | 71.3<br>(10.5) | 2.26* | .03 |
| 4 | 12.5<br>(8.6) | 23.0<br>(9.8) | 3.54** | .001 | 72.1<br>(13.0) | 66.0<br>(9.4) | 1.71 | .09 |
| 5 | 15.5<br>(11.3) | 27.3<br>(11.0) | 3.29** | .002 | 63.6<br>(15.4) | 55.0<br>(9.7) | 2.13* | .04 |
| 6 | 15.3<br>(11.3) | 27.3<br>(11.4) | 3.31** | .002 | 63.6<br>(15.5) | 54.9<br>(10.0) | 2.12* | .04 |
| 7 | 10.0<br>(10.0) | 19.7<br>(11.9) | 2.74** | .009 | 60.0<br>(14.2) | 52.7<br>(10.1) | 1.87 | .07 |

LTM = low trait mindfulness; HTM = high trait mindfulness. Values for the independent-samples t-test (df = 37) and corresponding p-values are also reported. Significant differences were found across all solutions except k = 4 and k = 7 for State 2. \* $p < .05$ , \*\* $p < .01$

**Figure 2.**
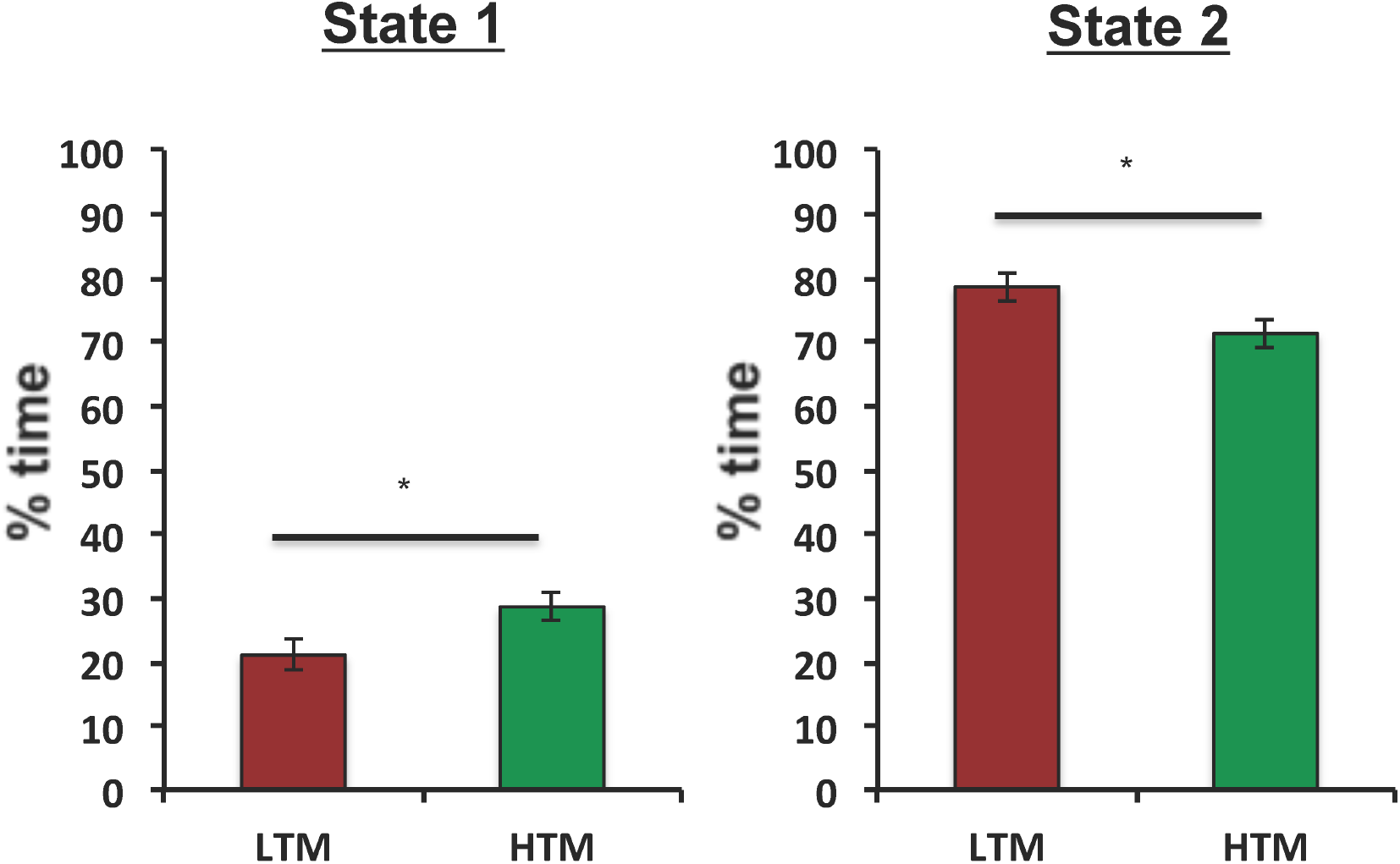
High trait mindfulness (HTM) individuals spend significantly greater time in a “task-ready” brain state (State 1), and significantly less time in an “idling” (State 2) state compared with low trait mindfulness (LTM) individuals. Results shown here are for k = 3 in k-means clustering. Error bars represent standard error of measurement. * *p* < .05

We tested the robustness of this finding in the higher cluster-number solutions, considering a state transition to be a change between any two states, not just into or out of State 1 (Table 2). The number of state transitions was significantly greater in the HTM group regardless of the number of clusters specified. Interestingly, as k increases, the average length of time spent in State 1 shows an increasingly larger difference between the HTM and LTM groups (HTM > LTM), reaching statistical significance when k = 7 (t_37_ = 2.31, p = .03) (Table 3).

**Table 2.** Mean and standard deviation of the number of state transitions for increasing values of k.

| Clusters | State transitions |  |  |  |
| --- | --- | --- | --- | --- |
| | LTM | HTM | $t$ | $p$ |
| 3 | 16.9 (7.6) | 22.5 (7.5) | 2.33* | .03 |
| 4 | 20.6 (7.7) | 27.2 (6.7) | 2.86** | .007 |
| 5 | 28.1 (8.5) | 33.8 (8.0) | 2.16* | .04 |
| 6 | 27.7 (8.9) | 33.7 (8.7) | 2.13* | .04 |
| 7 | 31.1 (8.0) | 38.2 (9.9) | 2.45* | .02 |

LTM = low trait mindfulness; HTM = high trait mindfulness. Values for the independent-samples $t$ -test ( $df = 37$ ) and corresponding $p$ -values are also reported. Significant differences were found across all values of $k$ . \* $p < .05$ , \*\* $p < .01$

**Table 3.**
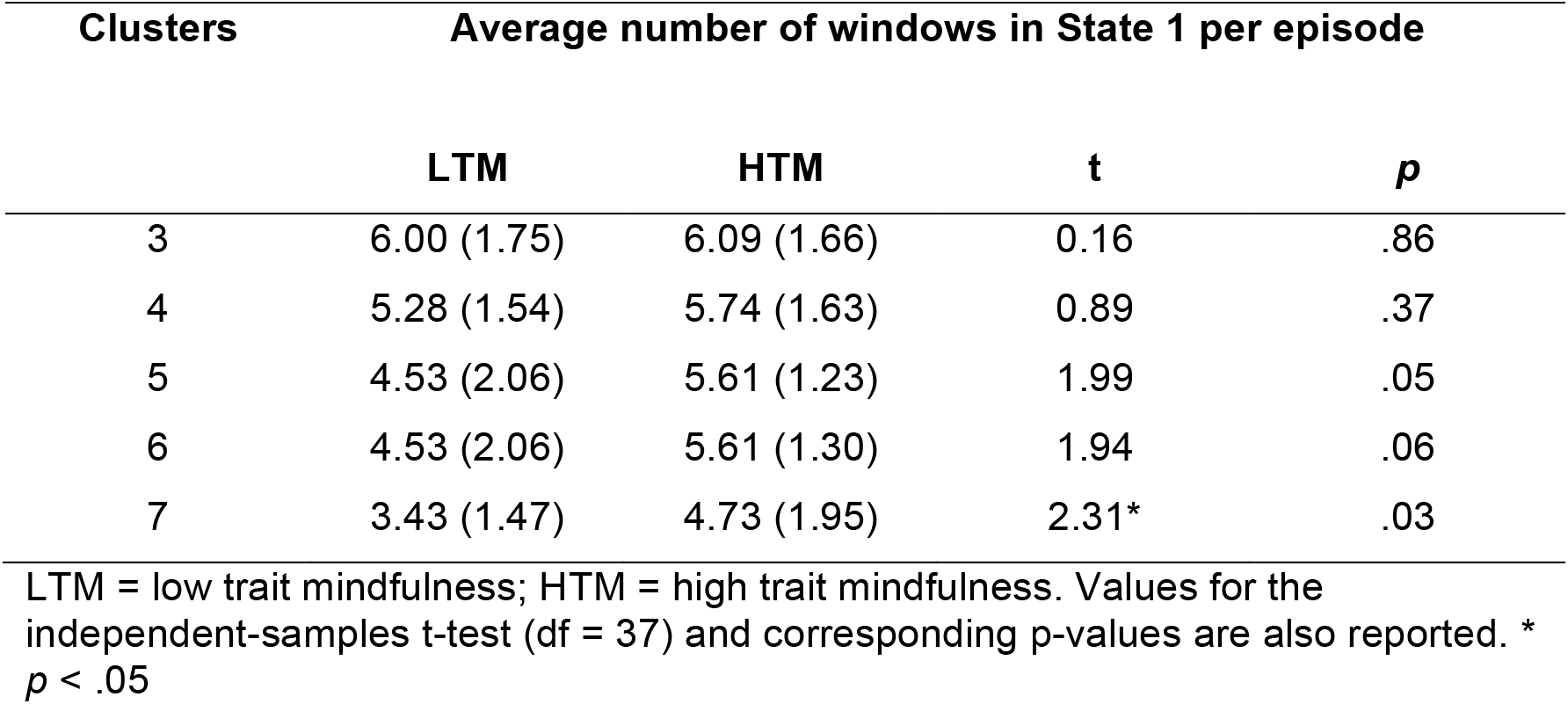
Mean and standard deviation of the average number of windows spent in each State 1 episode.

### 3.4 High trait mindfulness is associated with more state transitions

The greater time spent by HTM participants in State 1 might be due to two reasons. First, these participants might be transitioning more frequently into these states, but have an equivalent amount of dwell time in each State 1 episode. Alternatively, HTM participants might transition into State 1 the same number of times (or fewer) as LTM participants, but spend a longer dwell time in each State 1 episode. We tested this statistically, and found support for the first hypothesis. HTM individuals transitioned between states significantly more often than LTM individuals (t_37_ = 2.33, p = .03), but did not spend more time in State 1 per episode on average (t_37_ = 0.16, p = .86) (Figure 3).

**Figure 3.**
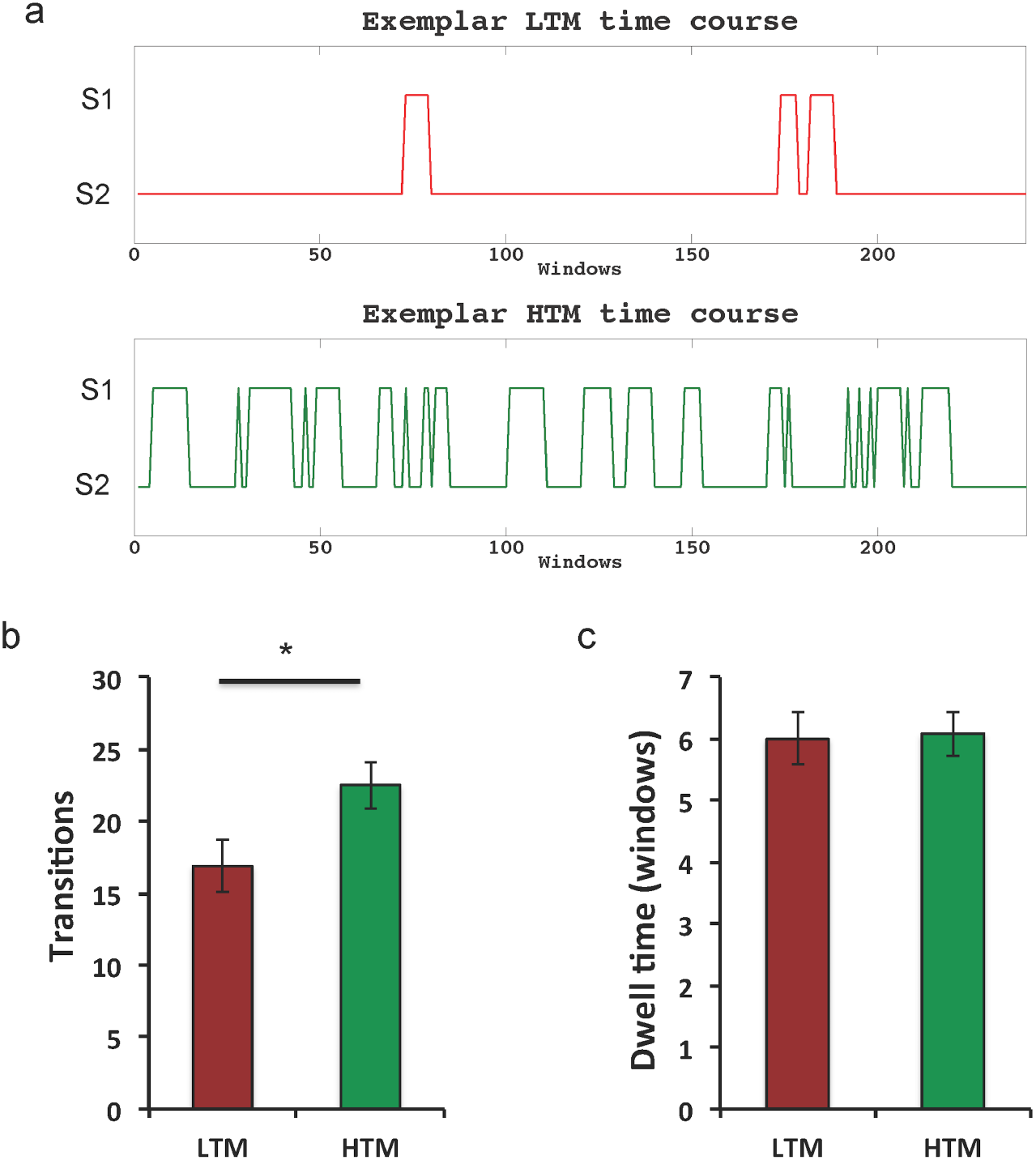
Dwell time and transitions in connectivity states. a) Time courses of involvement in each state (S1 and S2) during the resting-state scan for an exemplar high trait mindfulness (HTM) (subject 108) and low trait mindfulness (LTM) (subject 94) participant. b) On average, HTM individuals made more transitions between states, but c) did not spent a greater amount of time in each S1 episode. Error bars represent standard error of measurement.

### 3.5 Mindfulness or vigilance?

Previous work has suggested the two robustly observed polar DFC states are related to vigilance or sleepiness (Wang, et al., 2016). In the current dataset, the LTM group committed more lapses (Basner and Dinges, 2011) than the HTM group on the 20-minute PVT (11.7 vs. 5.2), and our previous dataset showed a significant correlation between BCT accuracy and PVT lapses (Wong et al., in press). To test whether vigilance or arousal might account for our observed effects, we repeated our comparison of the DFC variables between groups using PVT lapses as a covariate. In this updated model, the group difference in time spent in State 1 (F_1,36_ = 7.33, p = .01) and number of state transitions (F_1,36_ = 6,36, p = .02) remained significant, and was in fact slightly stronger than in the direct comparison.

### 3.6 Greater DMN and VAN connectivity in individuals with high trait mindfulness

We next generated static connectivity matrices of resting-state data by computing the average connectivity across all MTD windows for each subject (Figure 4a). Based on prior literature (Fox, et al., 2014; Young, et al., 2017), we selected four major networks that have been implicated in mindfulness and mindful attention – the default mode network (DMN), dorsal attention network (DAN), ventral attention/salience network (VAN), and the executive control network (ECN). Average within-network connectivity was computed for each of these networks, as well as the anti-correlation between DMN and each of the three task-positive networks. Results were FDR-adjusted (to yield q-values) across the seven planned comparisons. We found significantly greater within-network connectivity in the HTM group in the DMN (t_37_ = 2.37, p = .02, q = .047) and the VAN (t_37_ = 2.47, p = .02, q = .047), and significantly greater anti-correlations in the HTM group between DMN-DAN (t_37_ = 2.30, p = .03, q = .047) and DMN-VAN (t_37_ = 2.84, p = .007, q = .047). DMN-ECN connectivity did not differ significantly between the groups (t_37_ = 1.86, p = .07, q = .10) (Figure 4b).

**Figure 4.**
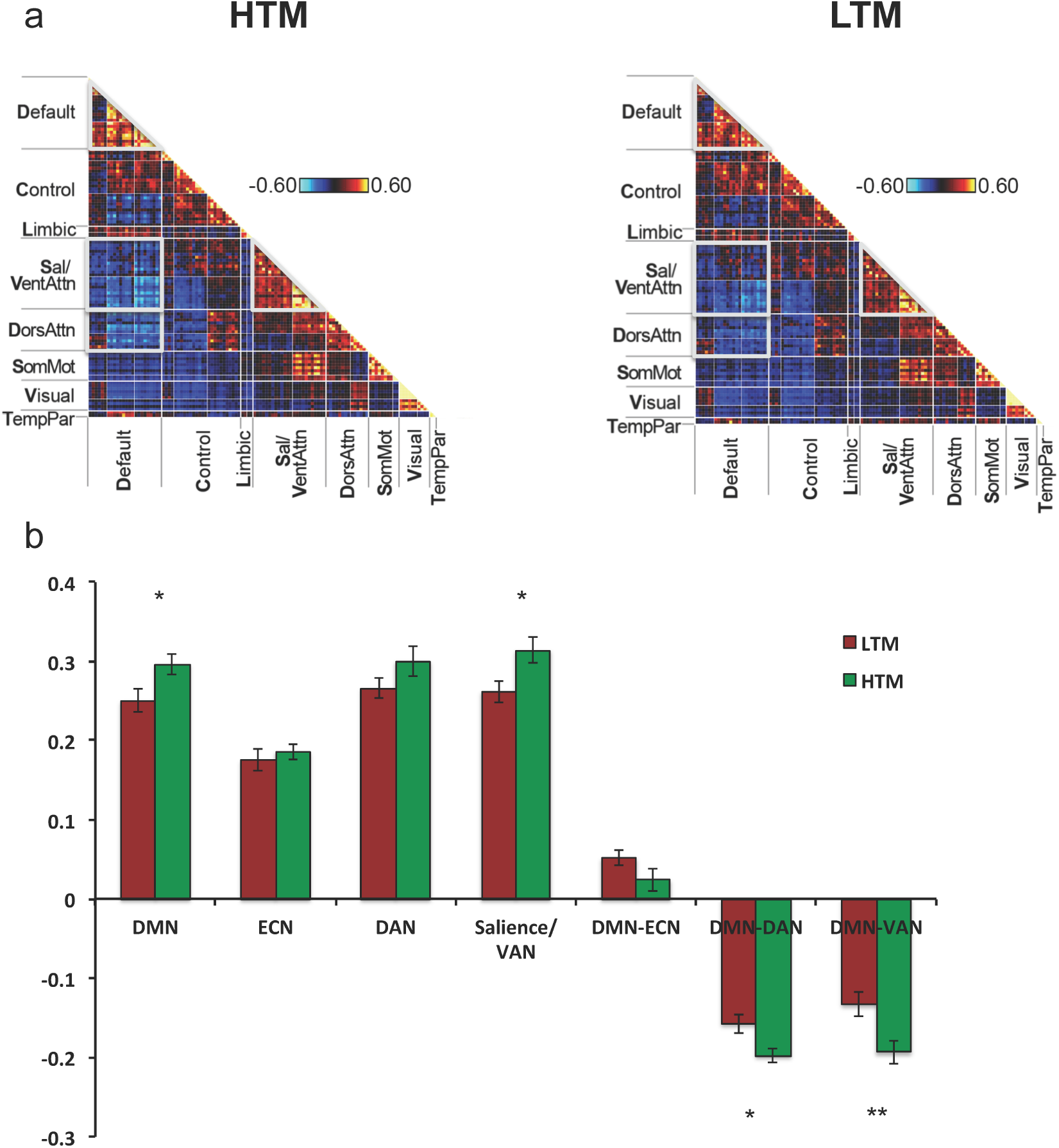
Static functional connectivity. a) Average resting-state static connectivity maps in the high trait mindfulness (HTM) and low trait mindfulness (LTM) groups. Units of connectivity are average coupling scores derived from the multiplication of temporal derivatives method. b) HTM individuals showed significantly stronger within-network connectivity in the DMN and salience/VAN networks, and significantly stronger anti-correlations between DMN-DAN and DMN-VAN. These significant comparisons are highlighted as areas bounded by the thick silver lines in panel a. All results remained statistically significant at q = .05 after correction using false discovery rate. Error bars represent standard error of measurement. Default/DMN = default mode network; Control/ECN = executive control network; DorsAttn/DAN = dorsal attention network; Sal/VentAttn/VAN = ventral attention network. * *p* < .05, ** *p* < .01.

### 3.6 Time spent in State 1 correlates with subjective mindfulness

Finally, we tested the relationship between our DFC states and subjective mindfulness as assessed by the Five-Facet Mindfulness Questionnaire (FFMQ) using Pearson correlations. Due to experimenter error, data from 3 participants were not available for analysis (2 HTM; 1 LTM). Neither the correlation between FFMQ and time spent in State 1 in the k = 3 solution (r = .32, p = .06; Figure 5) or time spent in State 2 (r = .32, p = .05) reached significance. However, the correlation with State 1 crossed the threshold of statistical significance when k = 4 and 7, whereas the correlation between FFMQ and State 2 was not significant for all k > 3 (Table 4). We also tested for correlations in the static connectivity metrics that differed between the HTM and LTM groups. FFMQ scores were significantly negatively correlated with DAN-DMN connectivity (r = -.37, p = .03), VAN-DMN connectivity (r = -.35, p = .03), and showed a trend-level correlation with connectivity within the DMN (r = .32, p = .06) (Figure 5). However, using FDR-adjustment, none of these correlations were statistically significant (all q = .061).

**Table 4.** Correlations between total score on the Five-Facet Mindfulness Questionnaire (FFMQ) and time spent in States 1 and 2 with increasing values of k.

| Clusters | State 1 |  | State 2 |  |
| --- | --- | --- | --- | --- |
| | $r$ | $p$ | $r$ | $p$ |
| 3 | .316 | .06 | -.324 | .05 |
| 4 | .348* | .04 | -.157 | .36 |
| 5 | .302 | .07 | -.170 | .32 |
| 6 | .306 | .07 | -.193 | .26 |
| 7 | .393* | .02 | -.144 | .40 |

\* $p < .05$

**Figure 5.**
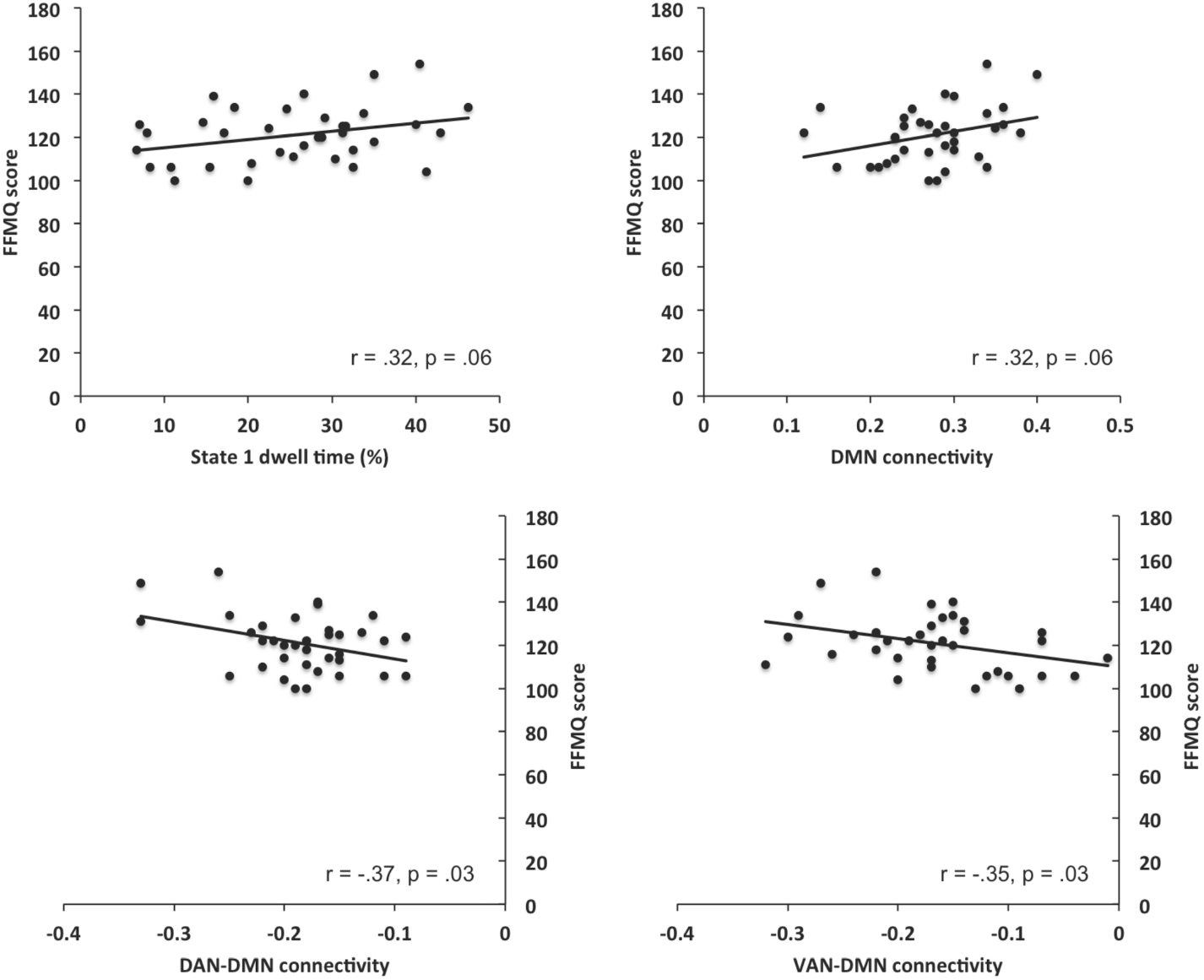
Correlations between subjectively reported mindfulness on the Five-Facet Mindfulness Questionnaire (FFMQ) and connectivity metrics. Top left: percentage of time spent in the task-ready state. Top right: connectivity within the default-mode network (DMN). Bottom left: connectivity between the dorsal attention network (DAN) and the DMN. Bottom right: connectivity between the ventral attention network (VAN) and the DMN. Using false discovery rate adjustment, none of these correlations were statistically significant (q = .061).

### 3.7 Potential confounds

Head motion is known to confound the effects of rs-fMRI data, even in datasets of relatively high quality (Power, et al., 2012). We tested whether there were systematic differences in the amount of head motion (framewise displacement) between the HTM and LTM group, and found that there were not (t_37_ = 0.84, p = .41), suggesting that this did not account for the group differences of interest.

As global signal was removed from the fMRI data prior to analysis, we investigated whether this variable might have differed between the HTM and LTM groups. Again, we found no significant difference between the two groups in global signal power (t_37_ = −0.07, p = .94).

## 4. Discussion

Mindfulness is a trait-like characteristic, as shown in this study by the good test-retest reliability of performance on a breath-counting task. Having confirmed this, we proceeded to investigate differences in resting-state functional connectivity between individuals high and low on trait mindfulness. Using a sensitive method of DFC analysis, we found that HTM individuals spend more time in a brain state characterized by high within-network connectivity, and between-network separation (State 1; the task-ready state (TRS)) due to the fact that they transition into this state more often than LTM individuals. Time spent in the TRS was associated with subjective reports of mindfulness. Static functional connectivity was also distinctly different between these groups, particularly for connections originating from the default mode network. Finally, we demonstrated that differences between the HTM and LTM group persisted after controlling for performance on a test of vigilance. Taken together, these results suggest that toggling between brain states in general, and into the TRS in particular, may be a neural substrate of the superior awareness of present-moment experience that is a feature of the mindful brain.

### 4.1 Interpreting DFC brain states

The first key finding of this study was that the HTM group spent significantly more time in a brain state with features suggesting greater brain segregation and integration. This result is in line with the findings of Mooneyham et al. (2017), who conducted DFC analysis on fMRI data collected during a period of mindful breathing. These authors found that the dwell time in a “focused attention” state correlated with dispositional mindfulness as measured by subjective report. Similarly, Marusak et al. (in press) report that trait mindfulness in children and adolescents was associated with less time spent in a state previously linked to mind wandering.

HTM participants also spent relatively less time in an “idling” state relative to LTM participants in the k = 3 solution; this is typically the predominant DFC state observed during resting-state scans, and one in which within-network connectivity is relatively weak (Allen, et al., 2014). Based on the topography of the idling state, it has been suggested that this state represents periods of lower information transfer (Tian, et al., 2018), perhaps to save on energetic costs when processing demands are low. We note that this finding was not as consistent across different clustering solutions, and was not statistically significant when considering a weighted average of effect sizes.

Previous studies (including Mooneyham et al. (2017)) have claimed that connectivity states homologous to the TRS represent periods of higher arousal and/or attention. For instance, an experiment using sleep deprivation (Wang, et al., 2016) has linked State 1/State 2 homologs to high and low arousal using spontaneous eyelid closures as a behavioral marker. As attentiveness is an important component of mindfulness (Brown and Ryan, 2003; Shapiro, et al., 2006), higher vigilance could partially explain why HTM individuals spend a greater proportion of time in the TRS.

At the same time, we caution against an overly narrow interpretation of the cognitive correlates of the TRS and dynamic brain states in general. Given that DFC states are commonly derived in an unsupervised fashion (using k-means clustering), it remains an open question as to whether they represent comparable configurations across studies, or whether subtle differences in the derived networks across studies relate them to different cognitive constructs. Recent evidence does suggest that dynamic brain states are robust and reproducible (Abrol et al., 2017), making it likely that the current results are not an idiosyncrasy of the clustering method or the population under study. In our dataset, the DFC differences between the groups persisted even after controlling for vigilance. Moreover, time spent in the TRS was correlated with FFMQ scores. These additional pieces of evidence suggest that the TRS is related to components of mindfulness other than arousal, possibly meta-awareness (Jankowski and Holas, 2014) or cognitive flexibility (Moore and Malinowski, 2009).

More generally, we suggest that DFC states do not have one-to-one mappings onto cognitive states. Instead, we propose that State 1 (and its homolog in other studies), might be more broadly seen as a state of greater readiness for task performance that is relatively more expensive to maintain, while State 2 may be a default or idling state in which cognitive resources are being conserved, perhaps because participants are not attending to the contents of their consciousness. This broader framework of task-readiness accommodates recent findings showing that more efficient or optimized brain connectivity at rest is associated with good performance on cognitive tasks, and higher intelligence (Schultz and Cole, 2016). It also provides an adequate description of the currently known behavioral correlates of these brain states without ascribing specific cognitive functions to them, and initializes a testable framework that future research might be based on.

### 4.2 State transitions may represent more frequent refocusing in the HTM group

We found that the HTM individuals make significantly more state transitions than the LTM individuals, and this result echoes a recent report by Marusak et al. (in press). Cycling through a repertoire of functional states is a unique feature of the conscious brain (Barttfeld, et al., 2015), and it has been argued that this metastability is necessary for flexible cognition and behavior (Deco, et al., 2017; Tognoli and Kelso, 2014). State transitions in the resting state putatively correspond with changes in cognitive mode, possibly due to shifts in one’s locus of attention (Kucyi, et al., 2017) or the generation of spontaneous thought (Kucyi and Davis, 2014; Smallwood and Schooler, 2006). However, few studies of DFC to date have explicitly interrogated the behavioral correlates of these transitions (see (Li, et al., 2017) and (Cabral, et al., 2017) for two exceptions).

An elemental component of mindfulness training is instructing students on how to attend to the contents of the mind, and recognize when the attention has strayed from a desired locus (Lutz, et al., 2008). Moreover, trait mindfulness is strongly associated with meta-awareness, or the objective, higher-level awareness of the flow of thoughts, feelings, and sensations (Jankowski and Holas, 2014). Based on this, we speculate that the greater number of transitions observed in the HTM group may represent an increased frequency of checking in on different aspects of the internal or external environment (Kucyi, 2017). In this formulation, transitions represent points in time when an individual moves to a different awareness set, allowing them to refocus their attention if necessary (i.e. if they are trying to accomplish a particular task). In future, evidence from task-based fMRI data could be used to investigate this claim.

### 4.3 DMN and VAN static connectivity are stronger in the HTM group

In agreement with the existing mindfulness literature, static functional connectivity analysis revealed significant differences in the HTM and LTM groups in within-network connectivity in the DMN and VAN, and anti-correlations between these two networks. Mindfulness meditation practice typically leads to increased inter-network connectivity in hub nodes of the DMN (Brewer, et al., 2011; Creswell, et al., 2016). High trait mindfulness has also been associated with weaker connectivity between the DMN and the thalamus (Wang, et al., 2014), and greater DMN connectivity with task-positive regions in the brain (Prakash, et al., 2013; Way, et al., 2010), including connectivity with the salience network specifically (Doll, et al., 2015; Kilpatrick, et al., 2011).

The current results strengthen our confidence that the DMN and the VAN/salience network act in concert to promote dispositional mindfulness. The DMN has been implicated in mind wandering (Mason, et al., 2007), and the generation of both task-unrelated and stimulus independent thought (Christoff, et al., 2009). Furthermore, higher trait mindfulness is associated with lower levels of mind wandering (Fountain-Zaragoza, et al., 2016; Mrazek, et al., 2012). The ability to switch away from mindlessness to task-readiness is putatively mediated by VAN control (Sridharan, et al., 2008). Menon and Uddin (2010) have proposed a model in which the salience network coordinates the interaction between externally directed attention (supported by the ECN), and internally directed attention (supported by the DMN). Greater static connectivity within and between the VAN and DMN may thus represent the superior ability of these modules to detect mind wandering and switch out of this state if necessary.

We note that most prior research has investigated connectomic changes association with mindfulness or meditation training, or increases in state mindfulness. In contrast, the present study extends our knowledge by showing that individual differences in functional connectivity can be observed even when no intervention or mindfulness induction is used as an experimental manipulation. While it has been previously argued that dispositional mindfulness may be a different construct than cultivated or induced mindfulness (Grossman, 2008), our data suggest that their instantiation in the connectome is highly similar.

### 4.4 Objective vs. subjective measurements of mindfulness

Most previous studies have employed self-report (e.g. the MAAS) as a measure of trait mindfulness; however, subjective measurement is vulnerable to various kinds of response bias, and may lack construct validity (Grossman, 2011; Sauer, et al., 2013). The current study is the first to use an objective measure – breath counting ability – to assay trait mindfulness using neuroimaging, and our results lend support to the reliability and validity of this test. Although we found correlations between self-reported mindfulness on the FFMQ and some of our static and dynamic connectivity metrics, these correlations did not survive more stringent correction for multiple comparisons. Thus, while our imaging findings are convergent with previous studies in the literature, there may be subtle differences in the neural correlates of mindfulness as measured by objective or subjective tests. An interesting area for future research might be to directly compare the connectomic features of individuals high on trait mindfulness selected using these two methods.

## 5. Conclusion

As the literature on mindfulness expands, more investigators are calling for research into biomarkers of this trait. Obtaining such markers will be critical to understand the neural instantiation of mindfulness. Importantly, they may also stand as potential indicators of improvement in mindfulness-based therapies. Here, we propose two such candidates: the mindful brain switches more often into a task-ready state, and spends more time in that state overall. These features may underlie the flexibility and greater degree of awareness that characterizes mindful individuals.

## Acknowledgments

This study was funded by a donation from the Far East Organization. We acknowledge the assistance of Kian Foong Wong and Zuriel Hassirim in data collection and analysis.

---

**Supplementary Table 1.**
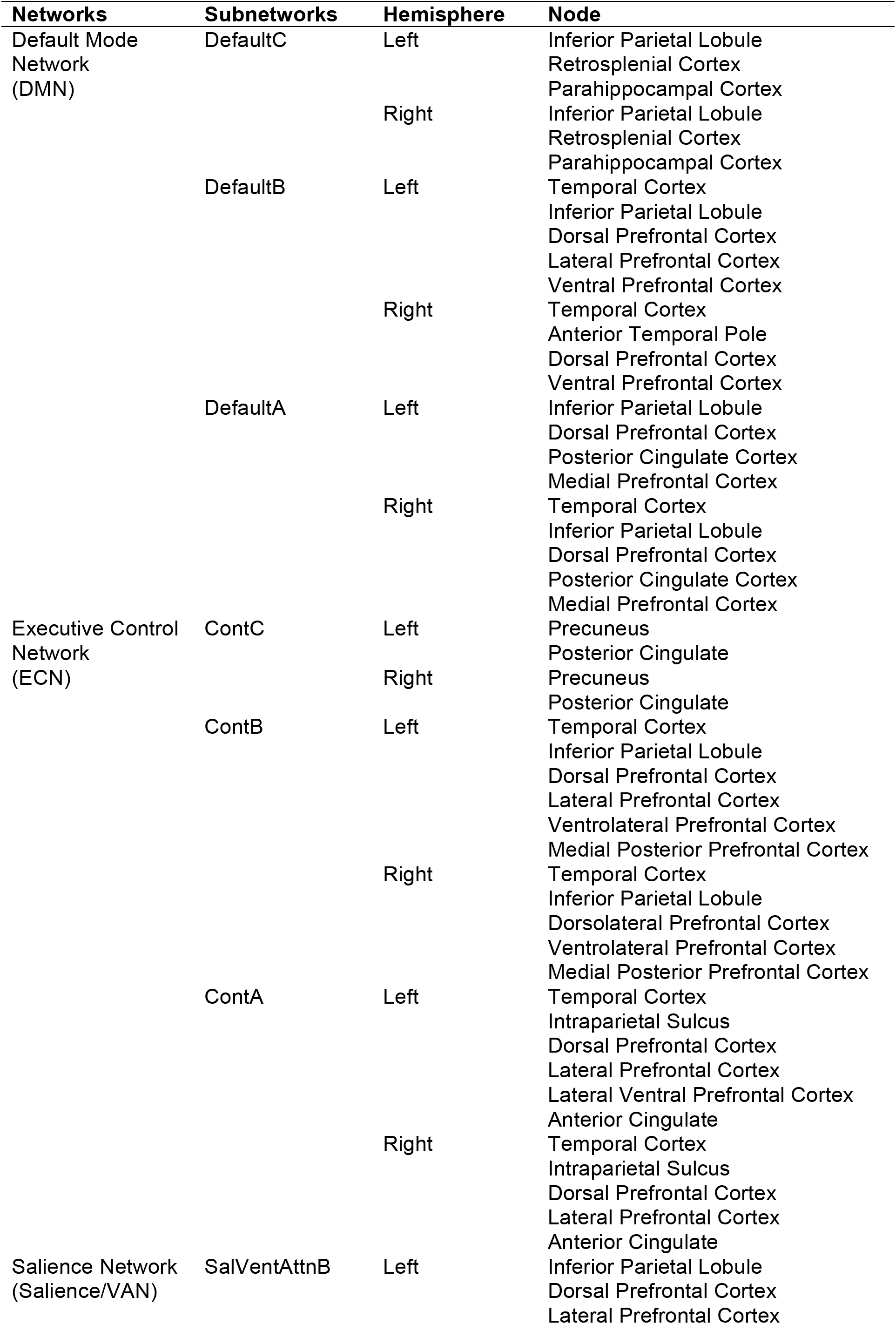
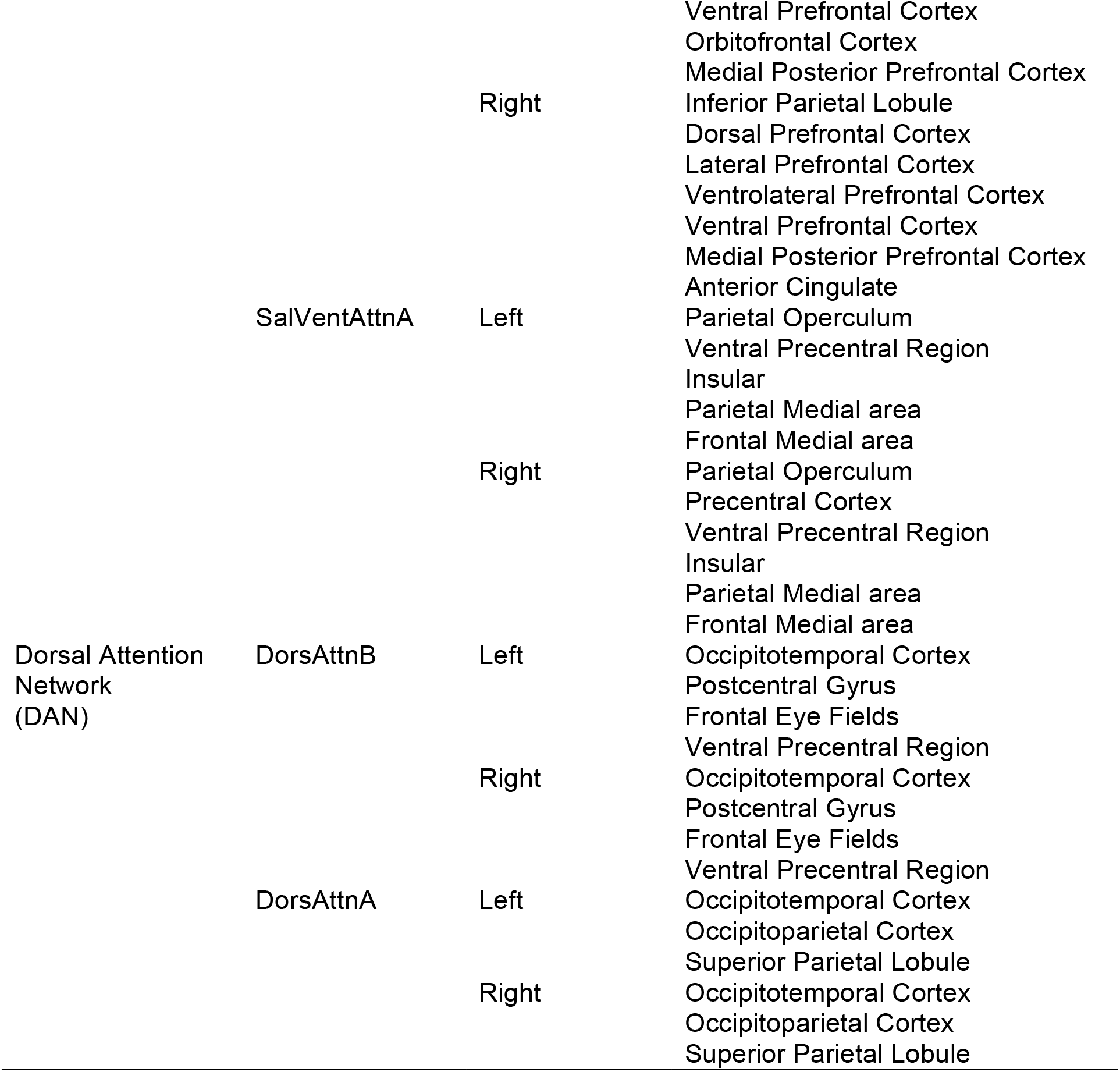
Names of individual network nodes.

